# DNA Double-Strand Break Movement in Heterochromatin Depends on the Histone Acetyltransferase dGcn5

**DOI:** 10.1101/2023.11.30.569406

**Authors:** Apfrida Kendek, Arianna Sandron, Jan-Paul Lambooij, Serafin U. Colmenares, Severina M. Pociunaite, Iris Gooijers, Lars de Groot, Gary H. Karpen, Aniek Janssen

**Affiliations:** Center for Molecular Medicine, University Medical Center Utrecht, Universiteitsweg 100, 3584 CG, Utrecht, the Netherlands; Department of Molecular and Cell Biology, University of California Berkeley, Berkeley, USA; Division of Biological Sciences and the Environment, Lawrence Berkeley National Laboratory, Berkeley, USA

## Abstract

Cells employ diverse strategies to repair double-strand breaks (DSBs), a dangerous form of DNA damage that threatens genome integrity. Eukaryotic nuclei consist of different chromatin environments, each displaying distinct molecular and biophysical properties that can significantly influence the DSB repair process. Specifically, DSBs arising in the compact and silenced heterochromatin domains have been found to move to the heterochromatin periphery in mouse and *Drosophila* to prevent aberrant recombination events. However, it is poorly understood how chromatin components, such as histone post-translational modifications, contribute to these DSB movements within heterochromatin. Using locus-specific DSB induction in *Drosophila* tissues and cultured cells, we identify that histone H3 lysine 9 acetylation (H3K9ac) is enriched at DSBs in heterochromatin but not euchromatin. We find that this enrichment is mediated by the histone acetyltransferase dGcn5, which rapidly localizes to heterochromatic DSBs. Moreover, we demonstrate that in the absence of dGcn5, heterochromatic DSBs display impaired recruitment of the SUMO E3 ligase Nse2/Qjt and fail to relocate to the heterochromatin periphery to complete repair. In summary, our results reveal a previously unidentified role for dGcn5 and H3K9ac in heterochromatin DSB repair and underscore the importance of differential chromatin responses at heterochromatic and euchromatic DSBs to promote safe repair.

## Introduction

DNA Double-Strand Breaks (DSBs), in which both strands of the DNA helix are severed, are one of the most harmful lesions known to occur in genomes. DSBs can be caused by both exogenous and endogenous sources, and when repaired improperly can directly result in the formation of aberrant chromosome structures linked to genetic diseases and cancer^1–4^. The two main DSB repair pathways in eukaryotic cells are Homologous Recombination (HR) and Non-Homologous End-Joining (NHEJ). HR requires 5’ to 3’ end-resection at the DSB site, which results in a single-stranded 3’ DNA end that invades and perfectly copies a homologous sequence to safely repair the DSB. In contrast, NHEJ involves simple ligation of the DSB, but often results in small nucleotide insertions or deletions at the repaired site^5,6^.

How different chromatin environments influence the cellular response to DSBs is less well understood^7^. Eukaryotic nuclei can be roughly subdivided into two types of chromatin: euchromatin, which is characterized by a more open conformation and contains most of the transcriptionally active protein-coding genes, and heterochromatin, which is more compact and relatively gene poor. Constitutive heterochromatin is one of the most prominent types of heterochromatin and is enriched at pericentromeric and subtelomeric regions. It contains highly repetitive DNA sequences, which are enriched for Histone H3 Lysine 9 di- and tri-methylation (H3K9me2/3) and its epigenetic reader protein Heterochromatin Protein 1a (HP1a in *Drosophila*, HP1α in mammals)^8–13^. Moreover, constitutive heterochromatin often coalesces into one or a few cytologically distinct domains in interphase nuclei^14,15^, which represent biocondensates that form through the distinct biophysical properties of HP1^16,17^.

The highly repetitive sequence content within this compact domain makes constitutive heterochromatin especially vulnerable to erroneous recombination events between homologous sequences present on non-homologous chromosomes. Previous work in *Drosophila* and mouse has revealed that heterochromatic DSBs can undergo end-resection within the heterochromatin domain, but only complete HR once relocated to the heterochromatin- or nuclear-periphery^18–21^. This DSB movement is thought to prevent aberrant HR repair within the repeat-rich heterochromatin domain^18,22^.

Several proteins have been found to be rapidly recruited to heterochromatic DSBs and promote DSB movement and HR repair. These include the structural maintenance of chromosomes 5/6 (SMC5/6) complex and the associated SUMO E3 ligases/Nse2 homologs, Cervantes (Cerv) and Quijote (Qjt)^18,19,23^. SMC5/6, as well as the other two SMC complexes cohesin and condensin, are conserved throughout the kingdoms of life^24^. The characteristic ring shape of SMC complexes enables them to bring multiple DNA sequences into close proximity and thereby play a crucial role in genome folding and organization^24–26^. SMC5/6 is known to promote genome stability in various pathways including chromosome segregation ^27,28^, telomere maintenance^29^ and DNA damage repair^30–33^. More specifically, the recruitment of SMC5/6 and its SUMO E3 ligase subunit Nse2 to heterochromatic DSBs in *Drosophila* is thought to mediate the SUMOylation of target proteins and in turn facilitate heterochromatic DSB movement through downstream signaling cascades^19,23^.

Although many histone modifications have been identified at euchromatic DSB sites^34,35^, histone changes at heterochromatic DSBs remain poorly understood. However, the distinct molecular and biophysical properties of heterochromatin suggest that DSB repair in this compact region requires specialized chromatin changes to allow DSB movement and repair protein recruitment. In line with this, we previously identified a role for the histone demethylase *Drosophila* KDM4A (dKDM4A) in specifically promoting the demethylation of H3K9me2/3 and H3K56me2/3 at heterochromatic DSB sites, thereby supporting DSB movement and repair pathway choice within heterochromatin^36,37^. Whether this demethylation is accompanied by additional changes in heterochromatin composition remains untested.

In budding yeast and mammalian cells, the acetylation of histone H3 Lysine 9 (H3K9ac) by the histone acetyltransferase General Control Non-repressed protein 5 (GCN5, dGcn5 in *Drosophila*) has been described to be involved in several types of DNA damage repair such as UV-damage^38^, oxidative damage^39^, and DSBs^40,41^. While one study reported a decrease^39^, the other studies revealed increases in H3K9ac levels in cells following damage induction^38,40,41^. However, these studies did not distinguish the responses in different pre-existing chromatin environments. It thus remains to be tested whether different chromatin environments, in particular the dense constitutive heterochromatin domain, differentially depend on H3K9ac for repair of damaged sites.

Using irradiation of *Drosophila* tissues and cells in culture, as well as our previously established *in vivo* single-DSB (DR-*white*) system in *Drosophila* tissues^20^, we here identify a specific increase in H3K9ac at heterochromatic DSBs. This H3K9ac is dependent on the histone acetyl transferase dGcn5, which is rapidly recruited to DSBs within the heterochromatin domain. Moreover, loss of dGcn5 results in defective DSB movement, as well as impaired recruitment of the SUMO E3 ligase Nse2/Qjt to heterochromatic DSBs. Together, our results identify a previously unrecognized role for dGcn5 and H3K9ac in the movement and repair of DSBs within the compact heterochromatin domain.

## Results

### The histone acetyl transferase dGcn5 deposits new H3K9ac marks at heterochromatic Double-Strand Breaks (DSBs)

In order to gain insights into the influence of chromatin factors on the process of DSB repair in heterochromatin, we sought to determine changes in chromatin composition associated with heterochromatic DSBs. We previously observed demethylation of histone H3 Lysine 9 di- and tri-methylation (H3K9me2/me3) at heterochromatic DSBs ^37^. We hypothesized that additional chromatin changes, such as histone acetylation, could be required to overcome the compact heterochromatic state and promote DSB movement and repair. To test this, we employed our previously established DR-*white* single DSB system in *Drosophila* (**Fig. 1A**)^20^. In this system, the DR-*white* construct is integrated in a single euchromatic or heterochromatic locus in the fly genome and contains an 18 base-pair recognition sequence targeted for DSB induction by the I-SceI endonuclease. To induce the expression of I-SceI and the consequent formation of DSBs, we heat-shocked third instar larvae containing eu- or hetero-chromatic DR-*white* insertions as well as a heat-shock inducible I-SceI (hsp.I-SceI) transgene. ChIP-qPCR (Chromatin Immuno-precipitation (ChIP) followed by quantitative PCR (qPCR)) was performed on chromatin extracts upon I-SceI-dependent DSB induction (**Fig. 1B**). Interestingly, we observed that the levels of the histone mark Histone H3 Lysine 9 acetylation (H3K9ac) increased at DSBs in three of the five heterochromatic DR-*white* lines (1.6 - 2.8-fold increase), whereas H3K9ac levels decreased or remained unchanged at three euchromatic insertion sites (**Fig. 1C, S1A**). Importantly, internal controls demonstrate that the antibodies used in these experiments selectively bind H3K9ac nucleosomes (**Fig. S1B, C**).

**Figure 1.**
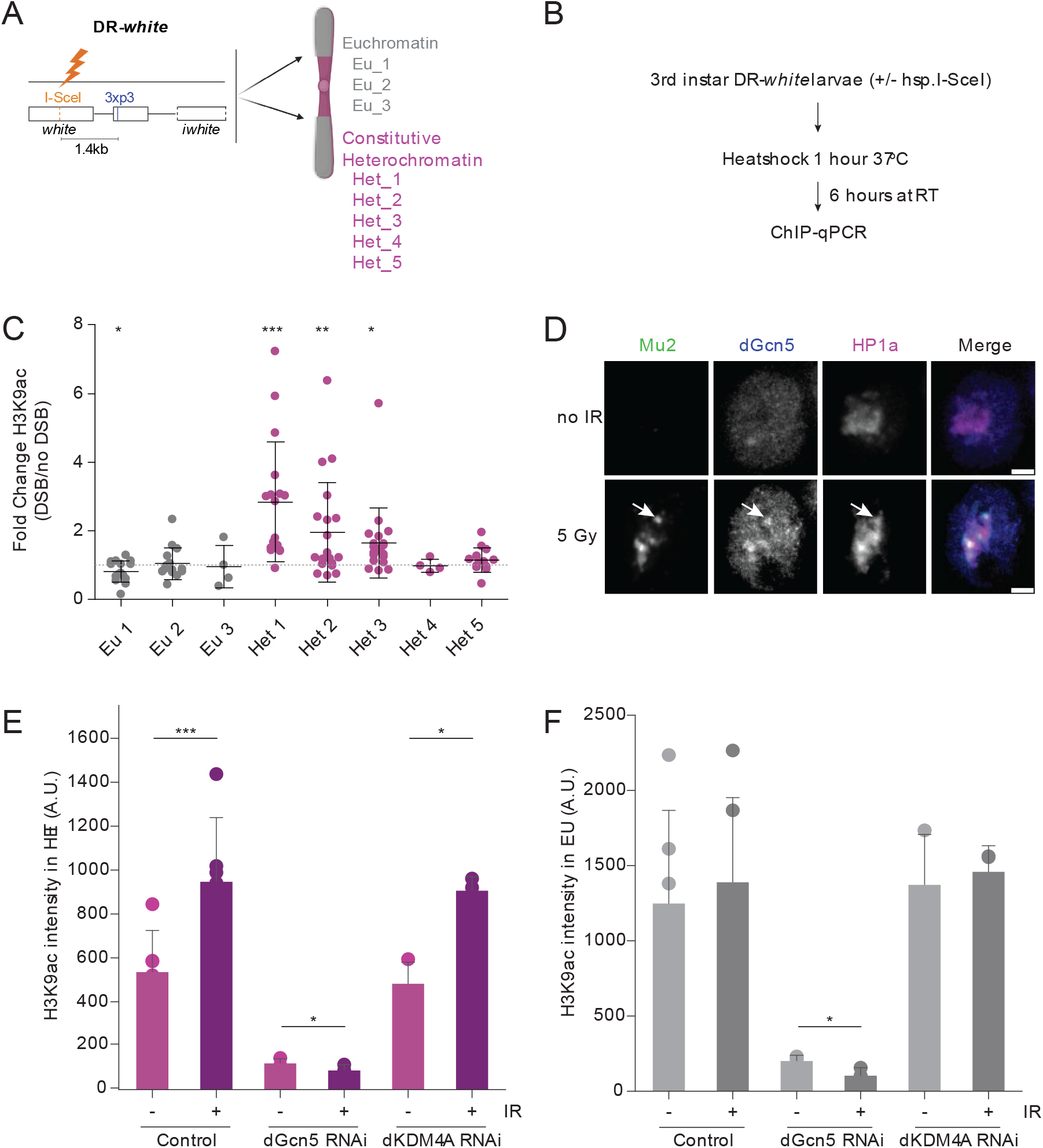
H3K9ac is increased at heterochromatic DSBs in a dGcn5-dependent manner. **A**. Schematic representation of the DR-*white* system in *Drosophila*. The DR-*white* construct is integrated in a single euchromatic or heterochromatic locus in the fly genome to generate different lines (Eu_1-3 and Het_1-5). The DR-*white* construct contains an 18-basepair DNA sequence targeted by the I-SceI endonuclease, resulting in the formation of a single DSB^20^. **B**. Scheme detailing the DSB induction protocol in DR-*white* fly lines. Chromatin was extracted 6 hours after DSB induction. DSBs were induced by a 1-hour heat shock (37°C) of third instar DR-*white* larvae containing a heat-shock promoter-driven I-SceI transgene (hsp.I-SceI). To control for heat-shock effects, all conditions (without/with hsp.I-SceI) are heat shocked. **C**. ChIP-qPCR analysis of H3K9ac levels at indicated DR-*white* loci. The qPCR primers reside 1.4 kb downstream of the I-SceI cut site (“3xp3” in *Fig. 1A*). Fold change is calculated by dividing the H3K9ac enrichment levels in the damaged (+ DSB, + I-SceI) samples by those in the controls (-DSB, no I-SceI). The grey dotted line represents a fold change of 1, in which H3K9ac levels are unchanged upon DSB induction. All qPCR results are relative to an internal control region (*yellow*), which is consistently enriched for H3K9ac. Error bars represent mean±SD from ≥4 independent experiments. **D**. Representative time-lapse images of non-irradiated (no IR) and irradiated (5 minutes after 5 Gy gamma-irradiation) Kc cells with fluorescently tagged Mu2 (green, DSB marker), dGcn5 (blue) and HP1a (magenta, heterochromatin marker). Arrows indicate dGcn5 localizing to a heterochromatic DSB. Scale bars = 2μm. **E, F**. Quantification of H3K9ac intensity levels in heterochromatin (*E*, H3K9me3-enriched, immunofluorescence stainings performed as in *Fig. S1I*) and euchromatin (*F*, low H3K9me3, see *Fig. S1I*) in non-irradiated versus irradiated control (yellow dsRNA), dGcn5-depleted and dKDM4A-depleted Kc cells. Irradiated cells were fixed 5-10 minutes after damage induction. Error bars represent mean +SD from ≥3 independent experiments. (*) P-value≤0.05, (**) P-value≤0.01 (***) P-value≤0.001, paired t-test. If not shown, P-value not significant (>0.05).

Previous work by others in mammalian cells as well as budding yeast have reported both the accumulation and the reduction of H3K9ac levels at DNA damage sites^38–41^. However, our ChIP-qPCR results indicate that increases in this histone mark are specific to DSBs in heterochromatin. This suggests that H3K9ac is differentially regulated in response to DSBs depending on the pre-existing chromatin domain.

It has been reported that H3K9ac deposition in *Drosophila* mainly depends on the histone acetyl transferase (HAT) dGcn5 and to a lesser extent on Elp3^42–45^. To assess which of these two HATs could be responsible for the acetylation of lysine 9 on histone H3 in *Drosophila*, we knocked down both proteins via RNA interference in *Drosophila* Kc cells (**Fig. S1D, E**), followed by immunofluorescence staining for H3K9ac. We find that depletion of dGcn5, but not Elp3, leads to decreased nuclear H3K9ac levels in undamaged cells, supporting the notion that dGcn5 represents the major acetyltransferase for H3K9ac in *Drosophila* (**Fig. S1F, G**)^46^.

Next, to test whether dGcn5 or Elp3 are recruited to heterochromatic DSBs, we fluorescently tagged these proteins in *Drosophila* cells in culture. We combined their expression with fluorescently tagged HP1a (Heterochromatin Protein 1a) to visualize heterochromatin and Mu2 (binds phosphorylated H2Av (γH2Av), MDC1 homolog) to visualize DSBs. Strikingly, we observed co-localization of dGcn5 with DSBs (Mu2) in the HP1a domain within five minutes following 5Gy irradiation (**Fig. 1D**), indicating that this HAT can be directly recruited to DSBs within heterochromatin. Consistent with previous observations, Elp3 localizes to the cytoplasm in undamaged cells^47^, and this pattern was not affected by the induction of DSBs via ionizing radiation (5Gy) (**Fig. S1H**). Together, this suggests that dGcn5, but not Elp3, localizes to heterochromatic DSBs and represents the main H3K9ac transferase in *Drosophila* cells. We therefore focused our subsequent analyses on the potential role of dGcn5 at heterochromatic DSBs.

The observed localization of dGcn5 to DSBs in heterochromatin, and the general role for dGcn5 in H3K9ac^42^ (**Fig. S1F, G**), led us to hypothesize that dGcn5 could be responsible for the observed increase in H3K9ac levels at heterochromatic DSBs (**Fig.1C**). Thus, we irradiated control and dGcn5-depleted cells and quantified the H3K9ac intensity levels in euchromatin (H3K9me3 – low) and heterochromatin (H3K9me3 - high) (**Fig. S1I**) five minutes after damage induction. In order to focus solely on DSB-dependent changes in chromatin composition, quantification was performed only on cells displaying at least one γH2Av focus (phosphorylated H2Av, used as a DSB marker) in the heterochromatin domain (**Fig. S1I, Fig. 1E**). In line with our ChIP-qPCR results at single DSBs *in vivo* (**Fig.1C**), we find a significant increase in heterochromatic H3K9ac levels after irradiation of cells in culture (**Fig. 1E**). In contrast, the H3K9ac increase was more modest and not significant in damaged euchromatic regions (**Fig. 1F**). This result further supports the idea that H3K9ac levels are differentially regulated in damaged euchromatin and heterochromatin. Importantly, depletion of dGcn5 abolished the damage-dependent increase in H3K9ac levels, suggesting that dGcn5 is required for the deposition of new H3K9ac marks at the sites of heterochromatic DSBs (**Fig. S1I, Fig. 1E, F**).

We previously found that the repair of heterochromatic DSBs depends on the removal of H3K9me2/3 by the demethylase dKDM4A^36,37^. Since unmethylated histone residues are chemically required to allow other types of epigenetic modifications, we hypothesized that dKDM4A-mediated demethylation is necessary for the deposition of H3K9ac at heterochromatic DSB. To test this, we irradiated control and dKDM4A-depleted cells and subsequently quantified the H3K9ac intensity levels at DSBs in euchromatin and heterochromatin (**Fig. S2A, Fig. 1E, F**)). Surprisingly, we found that the absence of dKDM4A did not affect the increase in H3K9ac levels at heterochromatic breaks. Consistent with our observations in cultured cells, we found that H3K9ac levels at heterochromatic DSBs were unaffected in irradiated wing discs of homozygous dKDM4A mutant larvae (ΔdKDM4A) (**Fig. S2B**)^37,48^. This suggests that the H3K9ac marks at DSBs in heterochromatin are not deposited on the same histones demethylated by dKDM4A and therefore indicates that H3K9 demethylation and acetylation are two independent events occurring at heterochromatic DSBs.

In conclusion, our data reveal that the histone acetyl transferase dGcn5 is rapidly recruited to heterochromatic DSBs and deposits new H3K9ac marks at the damaged sites. Moreover, this heterochromatin-specific increase in H3K9ac occurs independently of dKDM4A-mediated H3K9me2/3 removal.

### dGcn5 promotes timely movement and repair of heterochromatic DSBs

In *Drosophila*, heterochromatic DSBs relocate to the heterochromatin- and nuclear-periphery within thirty minutes after irradiation^18–20^. Since we observed a heterochromatin-specific increase in dGcn5-mediated H3K9 acetylation as early as five minutes after irradiation (**Fig. 1D-F**), we hypothesized that this local post-translational modification is one of the factors that promote heterochromatic DSB movement. To test this, we depleted dGcn5 in *Drosophila* cells in culture and assessed the presence of DSBs (γH2Av foci) in heterochromatin (DAPI bright^18,49^) and euchromatin (DAPI weak) at specific time points following 5Gy irradiation (**Fig. 2A**). Consistent with earlier findings^18^, we find in control cells that the number of heterochromatic DSBs peaked 10-20 minutes after irradiation (**Fig. 2B**). This peak was followed by a steep decrease in the number of γH2Av foci located within heterochromatin, indicative of DSB movement (**Fig. 2B**)^18^. Cells depleted of dGcn5 acquired the same number of heterochromatic DSBs within a similar timeframe as control cells (**Fig. 2B**). However, the number of DSBs remained high within the heterochromatin domain (30’ - 180’) and did not decline as steeply as in control cells (**Fig. 2A, B**). These results suggest that loss of dGcn5 results in delayed heterochromatic DSB movement when compared to the control. Additionally, in euchromatin (DAPI weak) we did not observe any differences in number of DSBs between control and dGcn5 RNAi cells (**Fig. 2A, C**).

**Figure 2.**
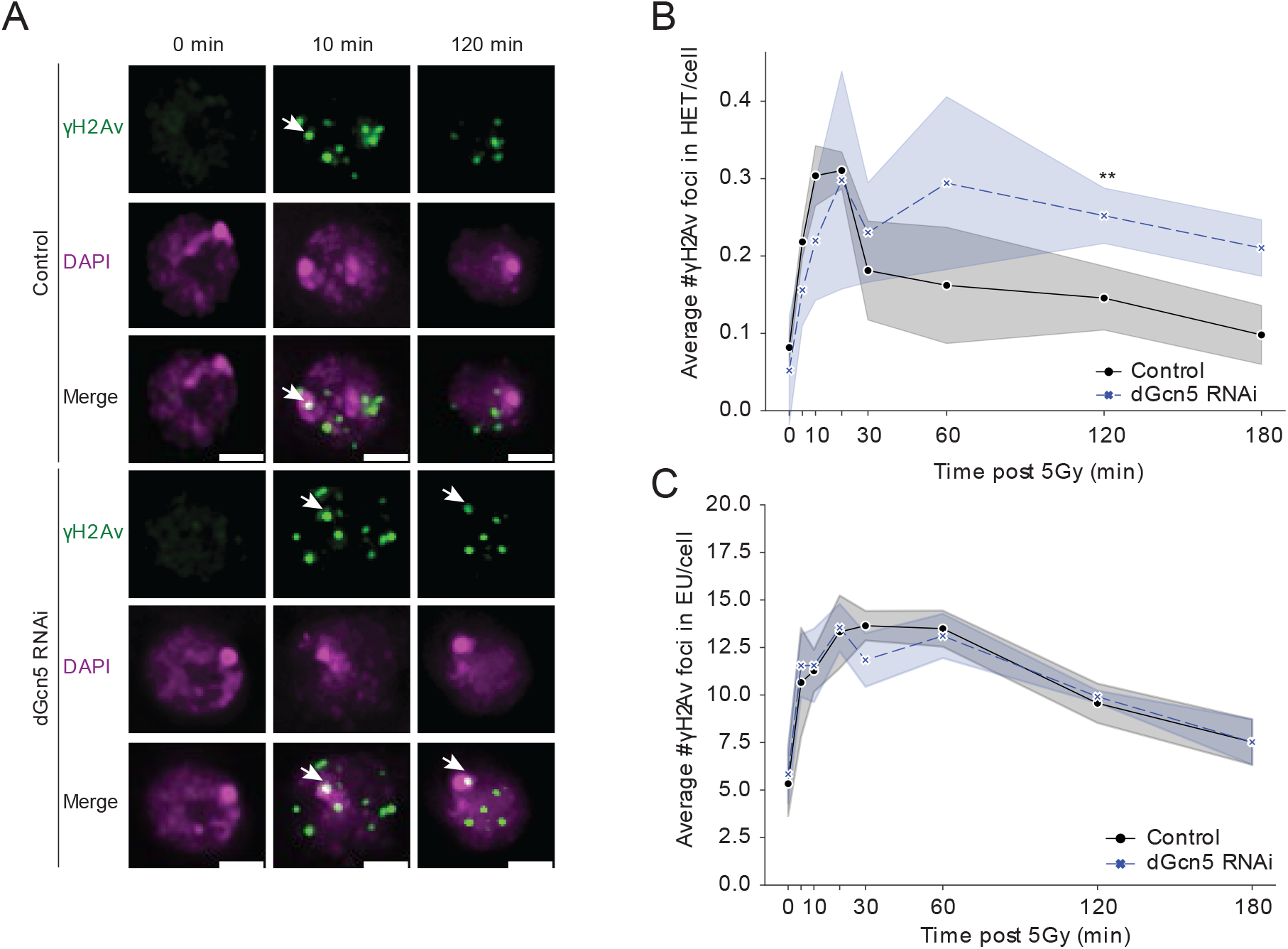
dGcn5 promotes repair of heterochromatic DSBs. **A**. Representative images of Kc cells depleted for yellow (control) and dGcn5 fixed at indicated time points following 5Gy irradiation. Cells were stained for γH2Av (green, DSB marker) and DAPI (magenta). White arrows indicate γH2Av focus in DAPI bright domain (heterochromatin). Cells at time point 0 were fixed without prior irradiation. Scale bar = 2μm. **B, C**. Quantification of images as in *A*. Average number of γH2Av foci in DAPI bright (*B*) and DAPI weak (*C*) is shown at indicated time points. Shade bars represent mean±SD from 3 independent experiments. (**) P-value ≤ 0.01, paired t-test. HET = heterochromatin, EU = euchromatin. If not shown, P-value not significant (>0.05).

Previous work found that dGcn5 interacts with components of the SWI/SNF complex in yeast and human cells to promote efficient H2A(X) phosphorylation at DSB sites^40,50^. In our system, we therefore expected to find a lower average number of γH2Av foci in DAPI weak^49^ (which represent ∼80% of the nuclear area, **Fig. 2A**) upon dGcn5 knockdown. However, we do not observe any differences in euchromatic γH2Av accumulation in control versus dGcn5-depleted cells, suggesting that either the remaining dGcn5 expression (**Fig. S1D**) is sufficient to fulfill the role in H2Av phosphorylation or perhaps a yet unknown mechanism can take over this role in *Drosophila*. Altogether, our data indicate that knockdown of dGcn5 specifically results in impaired heterochromatic DSB movement.

In previous studies, it was found that heterochromatic DSB movement is predominantly associated with repair by HR^18,21^. Early HR steps (e.g. end-resection) occur within heterochromatin, whereas the later steps of HR (i.e. Rad51 assembly) resume only after DSBs have moved outside of the heterochromatin domain ^18^. Since our data reveal a delayed heterochromatic DSB movement upon dGcn5 knockdown (**Fig. 2A, B**), we wished to address whether the early HR process within the heterochromatin domain is affected in the absence of dGcn5. ATR Interacting Protein (ATRIP) is one of the proteins involved in early HR repair by binding to the RPA-ssDNA complex^51^ and has been found to be efficiently recruited to both eu- and heterochromatic DSBs^18,51^. We performed live imaging of fluorescently tagged ATRIP within the heterochromatin domain (visualized with HP1a) following 5Gy irradiation in the absence and presence of dGcn5. By following ATRIP foci kinetics (time from appearance to disappearance) as well as its dynamics (movement) in the HP1a domain, we find that, in the absence of dGcn5, ATRIP foci in the HP1a domain display defective movement (**Fig. 3A-C**). In the control situation, ATRIP foci remained on average sixteen minutes in the HP1a domain (**Fig. 3C**), after which they either move (52%, **Fig. S3A**) or get resolved within the domain. In contrast, ATRIP foci in dGcn5 depleted cells remained twice as long inside the HP1a domain (on average forty-one minutes) before they moved out (32%, **Fig. S3A**) or resolve (**Fig. 3A-C**). This result is consistent with the observed retention of γH2Av foci within heterochromatin in irradiated fixed cells (**Fig. 2**). We do not find a defect in initial ATRIP recruitment to DSBs in heterochromatin (**Fig. S3B**), indicating that early end-resection steps can occur in the absence of dGcn5 and that specifically later steps (DSB movement and repair) are significantly delayed.

**Figure 3.**
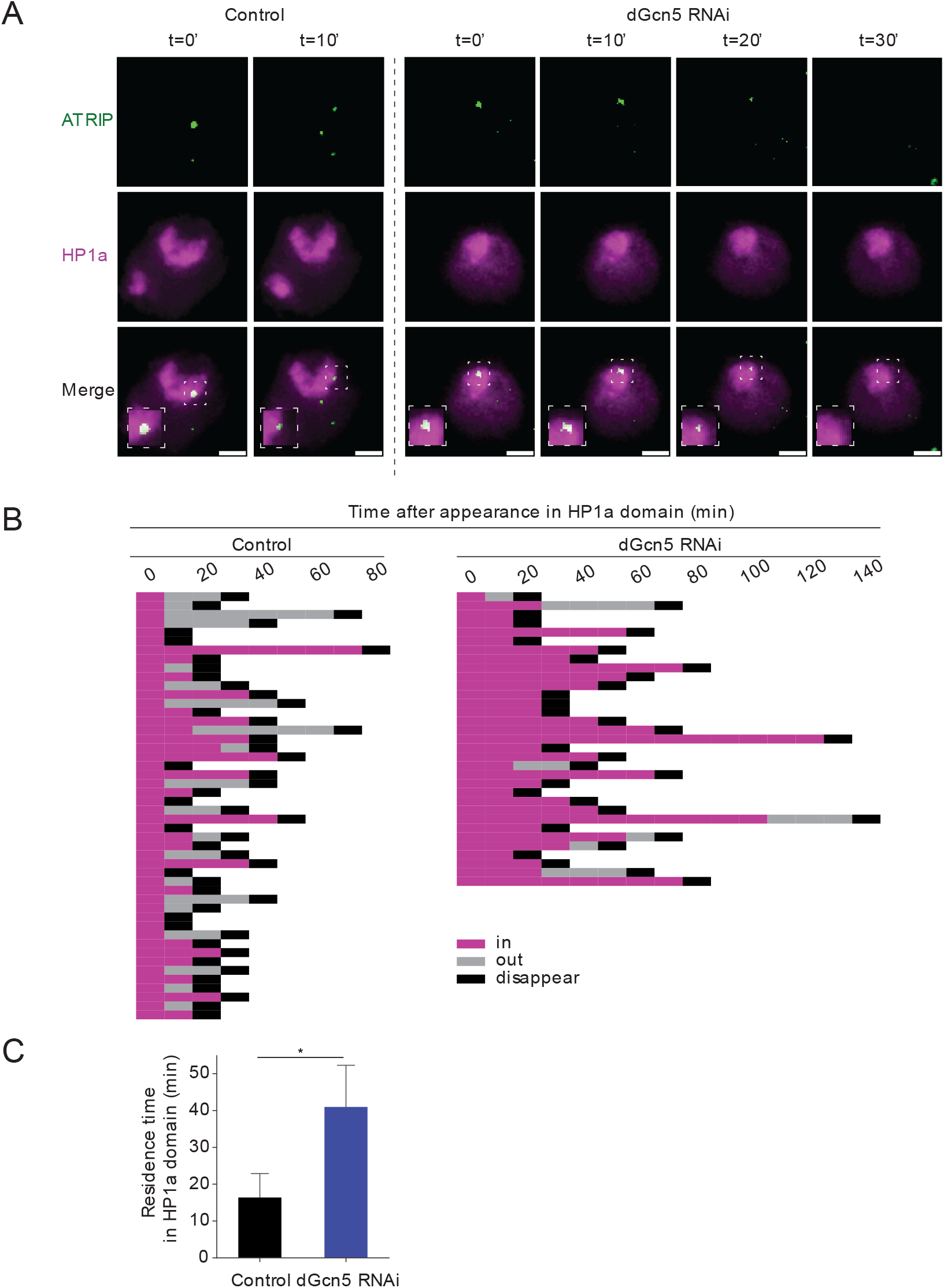
dGcn5 is required for DSB movement outside the HP1a domain. **A**. Representative time lapse images of ATRIP foci (green, HR protein) within the HP1a domain (magenta) in irradiated (5Gy) control (yellow dsRNA) or dGcn5 depleted Kc cells. Scale bars = 2μm. Insets are zoom-in views of heterochromatic ATRIP foci. **B**. Quantification of time-lapse movies as in *A*. ATRIP foci movement and kinetics plotted from their appearance in the HP1a domain (0’, magenta) to the timepoint they moved outside of the HP1a domain (out, grey) or resolved (disappear, black). Each row indicates quantification of one ATRIP focus. **C**. Quantification of the mean residence time of ATRIP foci within the HP1a domain. (*) P-value ≤ 0.05, paired t-test. Error bars represent mean +SD from 3 independent experiments.

To exclude that the observed DSB repair defects are due to changes in expression of DSB repair proteins, we performed bulk RNA-sequencing (RNA-seq) of dGcn5-depleted and control cells. These data reveal that the expression of known DSB repair proteins remained unchanged upon dGcn5 depletion (**Fig. S4A, Table S1-2**). dGcn5 knockdown also had no significant effect on the expression of cell-cycle regulated genes or cell-cycle progression based on RNA-seq data and EdU staining respectively (**Fig. S4A, B, Table S3**).

Together, these results suggest that H3K9 acetylation, mediated by dGcn5, plays a crucial role in promoting the movement and HR repair of heterochromatic DSBs.

### dGcn5 promotes the recruitment of the SUMO E3 ligase Nse2/Qjt to heterochromatic DSBs

Our results reveal that dGcn5 plays a role in promoting the movement of heterochromatic DSBs (**Fig. 2, 3**). We therefore sought to investigate the mechanism through which dGcn5 contributes to this process. One of the most important regulators of DSB movement in heterochromatin is *Drosophila* Nse2 (Cervantes/Quijote), part of the SMC5/6 complex^23^. In order to test the impact of loss of dGcn5 on *Drosophila* Nse2, we monitored the effect of dGcn5 depletion on the recruitment of Quijote (Qjt) to hetero- and eu-chromatic DSBs. We irradiated cells to induce the formation of DSBs and subsequently performed live imaging of fluorescently tagged Qjt and HP1a (**Fig. 4A-C**). In line with previous studies^19,23^, we find that Qjt overlaps with the complete heterochromatin domain (defined by HP1a) in non-irradiated cells, and that Qjt foci form in both eu- and hetero-chromatin upon irradiation (**Fig. 4A-C**). These previous studies also showed that Qjt foci remain in the damaged heterochromatic domain until ∼10 minutes after DSB induction, after which they move outside of the domain^19,23^. Interestingly, although we did not observe any difference in the appearance of γH2Av foci in heterochromatin 10 minutes after irradiation (**Fig. 2B**), we do find that dGcn5-depleted cells display a decreased number of Qjt foci inside the HP1a domain when compared to the control (**Fig. 4B**). Importantly, this phenotype was specific for Qjt recruitment to heterochromatic DSBs, as we observed no difference in the number of Qjt foci within euchromatin upon dGcn5 depletion (**Fig. 4C**). This result suggests that dGcn5 promotes the recruitment of Qjt to heterochromatic DSBs, which is subsequently required to guarantee the efficient and timely DSB movement outside the HP1a domain. Altogether, the DSB movement defects (**Fig. 2, 3**) and the impaired Qjt recruitment in dGcn5-depleted cells (**Fig. 4**) suggest that dGcn5, and the dGcn5-dependent increase in H3K9ac at heterochromatic DSBs, acts as a key player in the DSB movement pathway in heterochromatin by stimulating the recruitment of crucial regulatory factors, such as Qjt, to the damaged heterochromatin domain.

**Figure 4.**
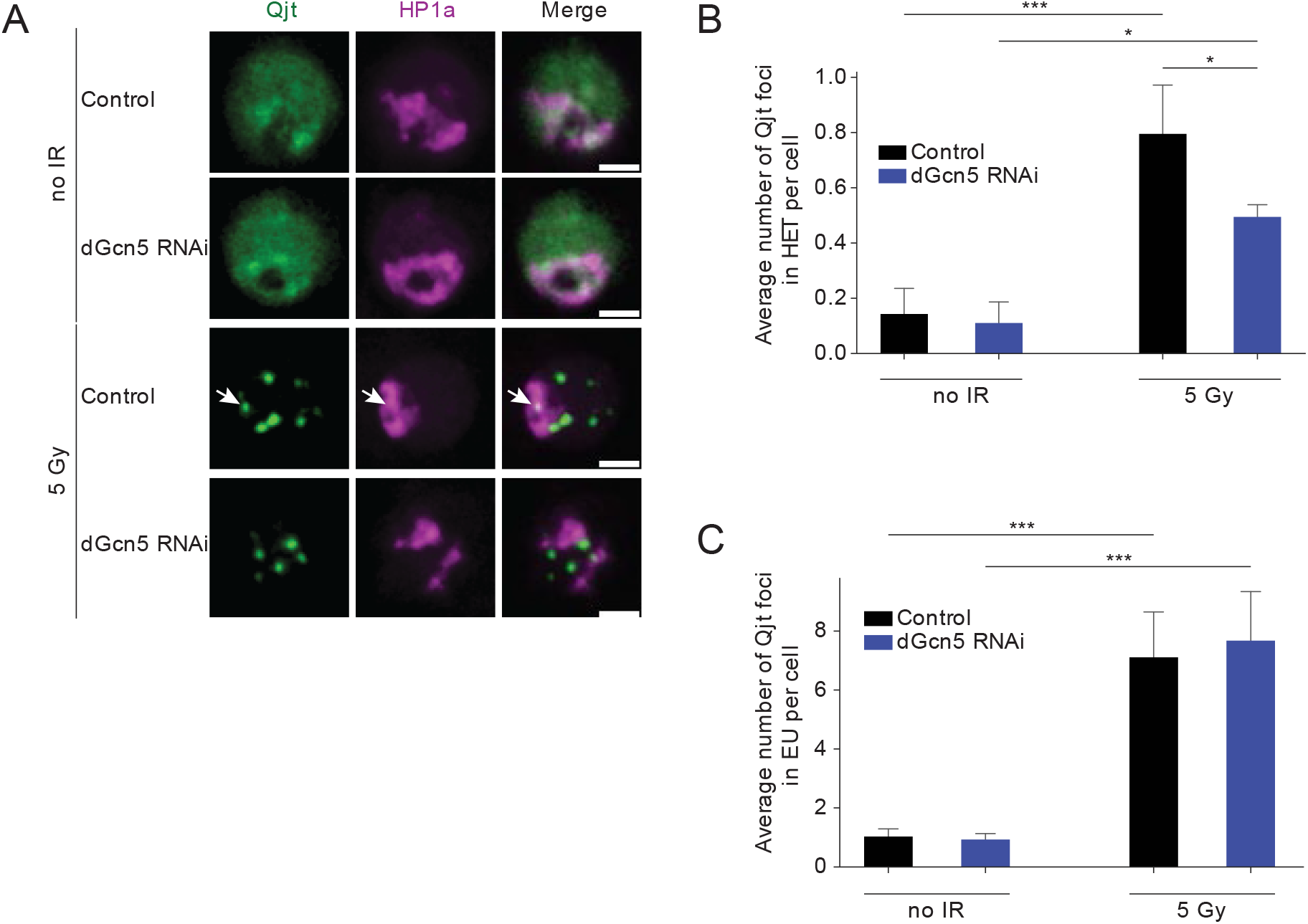
dGcn5 promotes Qjt recruitment at heterochromatic DSBs. **A**. Representative time-lapse images of Kc cells expressing HP1a (magenta, heterochromatin) and Qjt (Nse2, green) with indicated conditions. Arrowhead indicates Qjt focus residing inside the HP1a domain. Scale bars = 2μm. **B, C**. Quantification of Qjt foci inside the HP1a domain (*B*) and in euchromatin (*C*, outside HP1a domain) in non-irradiated versus 10 min after irradiation control (yellow dsRNA) and dGcn5-depleted Kc cells. HET = heterochromatin, EU = euchromatin. Error bars represent mean +SD from 3 independent experiments. (*) P-value≤0.05, (***) P-value≤0.001, one way ANOVA test followed by Tukey’s multiple comparison. If not shown, P-value not significant (>0.05).

### dGcn5 mutant flies depend on ATR for their survival

In order to confirm the contribution of dGcn5 to heterochromatic DSB repair *in vivo*, we employed dGcn5 heterozygous mutant flies that contain a premature stop codon on residue 333 (dGcn5[E333st]/+) (**Fig. 5A**)^44^. dGcn5 mutant flies are homozygous lethal but are heterozygous viable; heterozygous mutant larvae display a seventy percent reduction in dGcn5 mRNA and a significant decrease in H3K9ac levels (**Fig. S5A-C**)^44,52^. We dissected wing discs tissues from dGcn5[E333st]/+ third instar larvae and determined the number of DSBs (γH2Av foci) in heterochromatin (H3K9me3 region) before and after 5Gy irradiation (**Fig. 5B, C**). Consistent with our findings promotes repair of heteroci promotes repair of heterochhn cultured cells, we observed that, at 120 minutes after irradiation, DSBs accumulated in heterochromatin of dGcn5 mutant tissue, indicating that the movement of heterochromatic DSBs *in vivo* also requires dGcn5.

**Figure 5.**
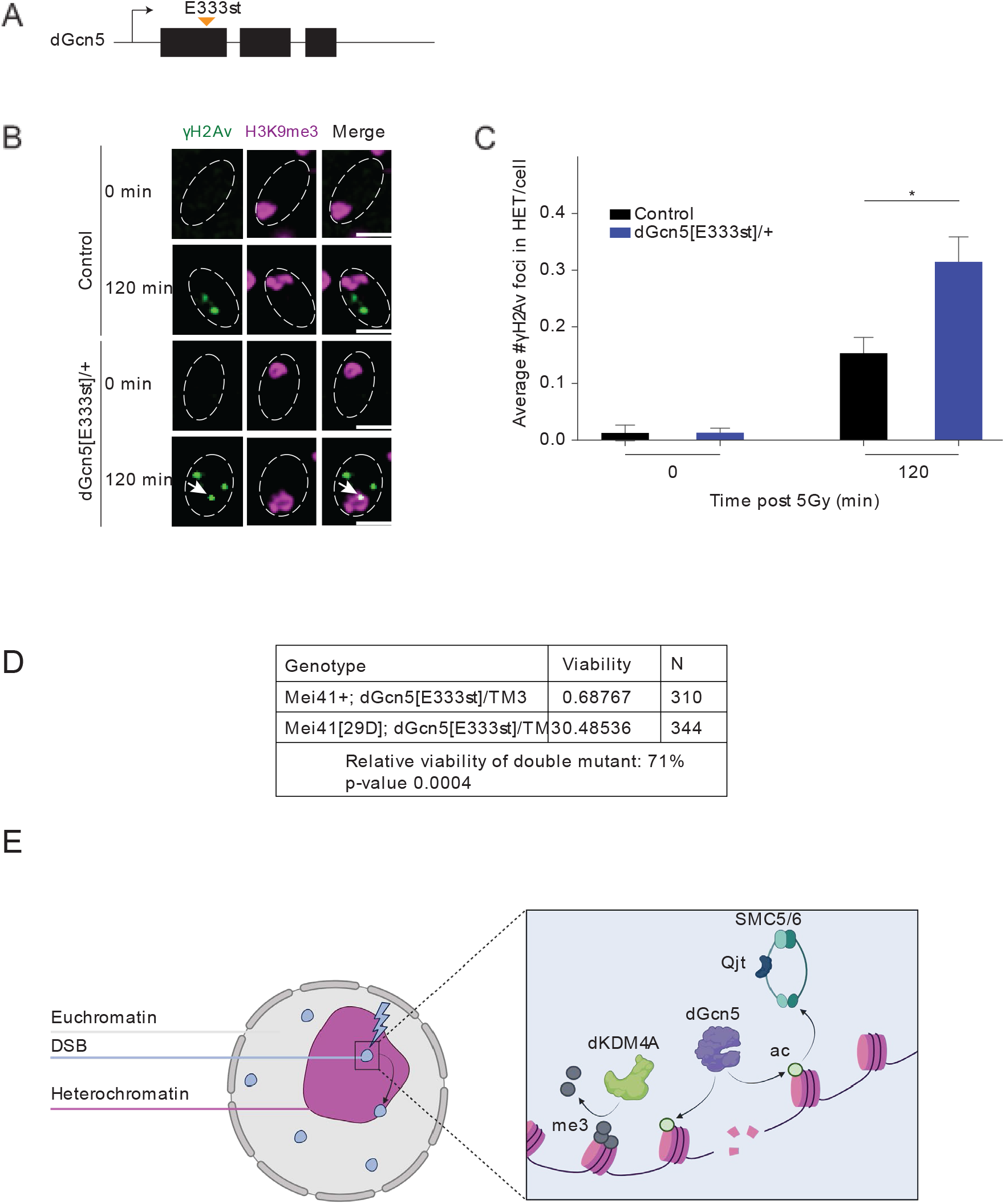
dGcn5 mutant is synthetically lethal with ATR mutant. **A**. Schematic representation of dGcn5[E333st] mutation. dGcn5 consists of 3 exons (black rectangles) and 2 introns (lines in between the black rectangles). The premature stop codon (E333st, orange arrow in first exon) is introduced in the three nucleotides encoding for the 333^rd^ amino acid^44^. **B**. Representative images of control and dGcn5[E333st]/+ wing disc cells, fixed without prior irradiation (0 min) or 120 min after 5Gy irradiation. Cells were stained for γH2Av (green, DSB marker) and H3K9me3 (magenta, heterochromatin marker). Dashed white lines indicate nucleus. Arrowheads indicate γH2Av inside the heterochromatic domain. Scale bar = 2μm. **C**. Quantification of images as in *B*. Average number of γH2Av foci in heterochromatin is shown at indicated time points. Error bars represent mean +SD from 3 independent experiments. (*) P-value≤0.05, one way ANOVA test followed by Tukey’s multiple comparison. If not shown, P-value not significant (>0.05). **D**. Synthetic lethality assay of dGcn5 mutant with ATR mutant. N=310 flies for the control line (dGcn5 mutant, wild-type ATR allele (mei41+; dGcn5[E333st]/+)) and n=344 flies for the cross between dGcn5- and ATR-mutant flies (mei41[29D]; dGcn5[E333st]/+). **E**. Model for the role of dGcn5-mediated H3K9ac in heterochromatic DSB repair. The local transfer of an acetyl group onto H3K9 by dGcn5 at heterochromatic break sites leads to recruitment of SMC5/6-Nse2 (Qjt) which in turn promotes movement and repair of heterochromatic DSBs. The deposition of new H3K9ac marks by dGcn5 is independent of dKDM4A-mediated demethylation of H3K9me3.

Finally, to assess the relevance of dGcn5-mediated H3K9 acetylation in living animals, we performed a synthetic lethality assay using dGcn5[E333st]/+ flies. The DNA damage repair kinase Ataxia telangiectasia and Rad3 related (ATR) is one of the proteins that rapidly respond to DSB events^53^. Moreover, it has been shown to promote heterochromatic DSB movement^18^. Combining dGcn5[E333st]/+ with an ATR truncation mutation (mei41[29D])^54^ results in a 30% decreased viability (**Fig. 5D**), highlighting the synergistic roles of dGcn5 and ATR in DSB repair. Altogether, these results suggest that dGcn5 mutant synthetic lethality with ATR is due to compromised heterochromatic DSB movement and repair.

## Discussion

Euchromatin and heterochromatin display different molecular and biophysical properties that can affect various aspects of DSB repair, including repair timing, DSB movement and repair pathway choice^22,35^. While in recent years research has started to uncover the impact of chromatin components on DSB repair, much still remains to be discovered about how epigenetic changes at DSBs are orchestrated in different pre-existing chromatin environments^22,35^. Since DSBs arising in heterochromatin specifically require movement towards the heterochromatin- or nuclear-periphery to avoid aberrant recombination events^18–20^, we hypothesized that this process is accompanied by unique changes in chromatin composition that ensure the efficient resolution of the damaged DNA. Here we tackled this question using our previously developed inducible single-DSB system in *Drosophila* tissues^20^, as well as irradiated *Drosophila* tissues and cultured cells. These results extend our previous findings on dKDM4A-mediated demethylation events at heterochromatic DSBs^37^ by revealing that timely repair of DSBs also depends on the dGcn5-dependent deposition of H3K9ac at the site of damage. In the absence of dGcn5, DSBs accumulate inside the HP1a domain and take twice as long to resolve or relocate outside heterochromatin (**Fig. 3**). Our data further suggest that defective DSB movement results from inefficient recruitment of the SMC5/6 complex component Nse2/Qjt to heterochromatic DSBs. We therefore propose a model in which dGcn5 mediates the local acetylation of heterochromatic H3K9 after DSB induction. Together with H3K9me2/3 demethylation, H3K9ac promotes movement and repair of the damaged DNA by facilitating the recruitment of Qjt to the HP1a domain (**Fig. 5E**).

The finding that H3K9ac levels increase almost exclusively at heterochromatic DSB sites constitutes novel information. Indeed, previous work has reported both the accumulation and the reduction of H3K9ac at DNA damage sites in human and yeast cells^38–41^. However, our *in vivo* ChIP-qPCR (**Fig. 1C**) and cell data (**Fig. 1E-F**) reveal that deposition of new H3K9ac marks is specifically enhanced at heterochromatic DSBs, suggesting that the levels of this mark are regulated according to the chromatin context in which the DSB is induced. This finding might therefore explain the conflicting results from previous studies in cells^38–41^, which focused on the general chromatin response to DSBs, without considering differences in chromatin composition prior to DSB induction. Altogether, our data reveal new insights into the specificities of DSB repair in heterochromatin and the importance of dGcn5 therein.

Interestingly, we were also able to observe dGcn5 localizing to DSB sites in heterochromatin, in line with previous *in vitro* and cell studies which reported dGcn5 being recruited to euchromatic DSBs and UV damage sites in both yeast and humans^38,40,41,55^. The finding that dGcn5 localizes to heterochromatic DSBs and specifically promotes their movement opens up new research directions. For example, how is dGcn5 exactly recruited to heterochromatic DSBs? Previously, we uncovered that movement and repair of heterochromatic DSBs in *Drosophila* depend on demethylation of H3K9me3 and H3K56me3 by the demethylase dKDM4A^36,37^. It would therefore be tempting to hypothesize that dGcn5 recruitment depends on dKDM4A, which would conveniently allow for control of H3K9ac deposition right after the demethylation of the same residue. However, in this study we find that the deposition of new H3K9ac marks by dGcn5 at heterochromatic DSBs does not require dKDM4A, suggesting that dGcn5 can be successfully recruited at break sites independently of dKDM4A. Several factors, including the MRN complex and the E2F1 transcription factor, have been previously implicated in promoting the localization of dGcn5 at euchromatic DNA damage sites in yeast and mammals (38, 40, 55). The recruitment of dGcn5 to heterochromatic DSBs might therefore depend on one of these proteins, although in this case another factor would have to confer specificity of the recruitment to heterochromatic DSBs.

We have shown that dGcn5 depletion leads to a decreased number of Qjt foci in damaged heterochromatin, suggesting that dGcn5 or H3K9ac promote the recruitment of Qjt to DSBs. It has been previously reported that Qjt is recruited to heterochromatic DSBs as part of the SMC5/6 complex^18,19,23^. Since none of the *Drosophila* SMC5/6 complex components possess a bromodomain (BRD, acetyl reader) (Uniprot database), we hypothesize that the newly deposited H3K9ac marks do not act as direct binding sites for SMC5/6-Cerv/Qjt at DSBs^56^. Nevertheless, one could envision a scenario in which Qjt is indirectly targeted by H3K9ac via an intermediate protein. Interestingly, a recent study in yeast proposed that dGcn5 and the SMC5/6 complex can directly interact, raising the possibility that the localization of Qjt at heterochromatic DSBs could also directly depend on dGcn5 binding^32^.

While the SMC5/6 complex is also enriched at euchromatic DSBs^31^, we did not observe a recruitment defect of Qjt to DSBs in euchromatin upon depletion of dGcn5 (**Fig. 4C**). This suggests that Qjt binding to DSBs in eu- and hetero-chromatin is regulated through different pathways, where heterochromatic DSB recruitment of Qjt is promoted by dGcn5-mediated H3K9ac, while in euchromatin other pathways are involved. Finally, we and others observed that Qjt and the SMC5/6 complex are already enriched in heterochromatin in undamaged conditions^18^ (**Fig. 4A**). While the role of the SMC5/6 complex in unperturbed heterochromatin is currently unknown, one intriguing possibility is that this localization pattern facilitates the fast recruitment of the complex to DSB sites upon deposition of new H3K9ac marks.

Another plausible role for H3K9ac at heterochromatic DSBs is the modulation of structural or biophysical properties of the damaged domain. dGcn5 has been reported to be important to promote chromatin accessibility at sites of DSBs and DNA replication^57,58^. Moreover, H3K9ac is associated with transcription and therefore highly enriched in euchromatin^59–61^. In the context of a condensate such as heterochromatin^16,17^, acetylation has been found to regulate the physical properties of chromatin by inhibiting phase separation^62^. Therefore, the enrichment of H3K9ac at heterochromatic DSBs could stimulate the formation of a more open (euchromatin-like) structure, in order to facilitate DSB movement, as well as the recruitment of repair factors.

Finally, the identified synthetic lethality between ATR and dGcn5 mutant flies suggests that the correct functioning of the dGcn5-mediated DSB movement pathway is important to preserve genome stability. Since both ATR^18^ and dGcn5 (this study) are required for proper movement of DSBs, we suggest that the decreased viability associated with the mutations in both proteins is due to the defective repair of heterochromatic DSBs. Although we cannot exclude that the synthetic lethality is caused by the contributions of dGcn5 to the recruitment of the SWI/SNF remodeling complex to DSBs^40,50^, we did not observe defects in the γH2Av accumulation and resolution in euchromatin (**Fig. 2C**), suggesting that euchromatic DSB repair can occur in the absence of dGcn5 in *Drosophila*.

Overall, our results provide an exciting illustration of how heterochromatin uniquely responds to a DSB, and modifies its chromatin features to promote successful repair. This is in line with the idea that chromatin components and chromatin-associated factors actively participate in the process of DSB repair and are fine-tuned depending on the respective chromatin domain^22,37,63^. Future research into how diverse chromatin environments affect and regulate the response to DSBs therefore represents a critical goal in order to understand not only how genomic stability is maintained across the eukaryotic nucleus, but also how failure to maintain genomic integrity can result in disease development.

## Materials and Methods

### Constructs

ATRIP, Qjt, dGcn5, elp3, Mu2 and HP1a were each cloned from cDNA generated from wild type Oregon R adult flies. cDNA for each gene was cloned into a pCopia plasmid backbone containing an N-terminal fluorescent tag (CFP, YFP, GFP or mCherry).

### Fly lines and genetic assays

All fly lines were reared at room temperature in standard medium unless otherwise stated. DR-*white* fly lines have been previously described^20^. Heterozygous dGcn5[E333st]/+ flies were obtained from the Bloomington *Drosophila* Stock Center (#9333) and the mei41[29D] mutant line was kindly provided by Tin Tin Su. The ΔdKDM4A [KG04636] deletion mutant was kindly provided by Mattias Mannervik^48^.

### Chromatin Immunoprecipitation-quantitative PCR (ChIP-qPCR)

Third instar larvae from DR-*white* fly lines containing the *hsp*.*I-SceI* transgene were heat-shocked at 37°C for 1 hour and snap-frozen in liquid nitrogen 6 hours after heat shock treatment. Larvae from DR-*white* fly lines not containing the *hsp*.*I-SceI* transgene were used as a control and subjected to the same heat shock treatment. Frozen larvae were stored at -80°C until chromatin extract preparation. For chromatin extract preparation, 80 larvae per line were homogenized, fixed and sonicated as described within the modENCODE project (https://www.encodeproject.org/documents/f890fde6-924c-4265-a60f-c5810401066d/, ChIP protocol by Kevin White lab)^64^. ChIP procedures were performed on chromatin extracts as previously described^65^. Each ChIP sample was prepared with 2μg of chromatin extract and 5μg of anti-H3K9ac antibody. qPCR was used to quantify the enrichment levels of histone marks of interest using the qPCR FastStart Universal SYBR Green Master Mix (Roche) and qPCR specific primers targeting the I-SceI site (3xp3) or the *yellow* gene as a control (Table S4).

### Cell culture, dsRNA production and transfection

S2 and Kc167 (Kc) cells were cultured at 27°C degrees in Schneider’s Insect Medium (Sigma-Aldrich) containing 10% FBS and CCM3 Serum-Free Medium (HyClone) respectively. For RNAi experiments, dsRNA was generated by first adding a T7-promoter sequence to the region of interest via PCR, followed by reverse transcription using the MEGAScript T7 transcription kit (Life Technologies) according to the manufacturer’s instructions (Table S4). 5-10μg of dsRNA was used to transfect 200.000 cells 5 days prior to irradiation. As a control, dsRNA targeting the *yellow* gene was always included. For live imaging experiments, cells were transfected with fluorescent proteins of interest approximately 6 hours prior to dsRNA transfection. For live imaging of Qjt, Kc cells were transfected with GFP-Qjt and Hygromycin resistance plasmids. Cells were selected for 2 weeks using 200μg/ml Hygromycin to generate Kc cells with stable GFP-Qjt expression. All transfections were performed using TransIT-2020 transfection reagent (Mirus) according to the supplier’s manual. Irradiation was performed by exposing cells to 5Gy of γ-rays in an IBL 437C machine.

### Reverse Transcription-quantitative PCR (RT-qPCR)

Knock down efficiency was verified by reverse transcription-quantitative polymerase chain reaction (RT-qPCR). Cells transfected with dsRNA were harvested after 5 days and RNA was extracted using the RNeasy Micro Kit (Qiagen) according to the manufacturer’s instructions. cDNA was obtained by using the iScript cDNA synthesis kit (BioRad). RT-qPCR reaction, using cDNA as template, was performed on a thermocycler using primers targeting *tubulin* as control. To test mRNA levels in dGcn5[E333st] larvae, RNA extraction and RT-qPCR were conducted as previously described^37^ (see Table S4 for primers used).

### Immunofluorescence (IF) and EdU staining

For IF staining on cells in culture, fixation was achieved by incubating cells with 4% Paraformaldehyde (PFA) in PBS for 5 minutes. Next, cells were incubated with 0.4% Triton-X diluted in PBS for 10 minutes to allow for permeabilization. Blocking was subsequentially performed by adding 5% milk, 0.4% Triton-X diluted in PBS for 1 hour. Primary antibodies were incubated overnight at 4°C, followed by secondary antibody and DAPI incubation for 2 hours at RT, all diluted in blocking solution. Finally, cells were mounted on glass slides using ProLong Diamond Antifade Mountant (Invitrogen).

For IF staining of tissue, wing discs were dissected from third instar larvae in 10μl of Schneider’s Insect Medium containing 10% FBS and fixed on a glass slide with 4% PFA for 5 minutes. Slides were dipped in liquid nitrogen and stored at -20°C in 99% ethanol. Slides were brought to RT, incubated in PBS for 20 min and subsequently blocked in 5% milk, 0.4% Triton-X diluted in PBS for 1.5 h. The following steps (primary and secondary antibodies incubation, mounting on glass slide) were performed as described above for fixed cells.

For EdU staining, Kc cells were incubated with 10μM EdU for 30 minutes. EdU detection was performed according to Click-iT EdU Imaging Kits Protocol (Invitrogen).

### Antibodies and dilutions

For ChIP-qPCR, experiments were performed with the anti-H3K9ac (Epicypher 13-0020, 13-0033) antibody. For IF experiments on tissues and cells, the following primary antibodies were used: anti-H3K9ac (rabbit, Epicypher 13-0033, 1:500), anti-γH2Av (mouse, Hybridoma Bank, UNC93-5.2.1, 1:250), FlexAble Coralite Plus 555 conjugated anti-H3K9me3 (rabbit, ab8898, 1:500). The following secondary antibodies were used: Alexa goat anti-rabbit 488 (1:600), Alexa goat anti-mouse 568 (1:600) (Invitrogen).

### Imaging and quantifications

Images were acquired using an RT DeltaVision microscope (DeltaVision Spectris; Applied Precision, LLC) using a 60X oil immersion objective (NA 1.40). For live imaging experiments, images were acquired every 10 minutes. Deconvolution was performed using SoftWoRx (Applied Precision, LLC) software. Image analysis was manually performed on deconvoluted images using the Fiji image analysis software.

### RNA-sequencing procedure

Kc cells were depleted of dGcn5 or yellow (control) for 5 days using dsRNA as described above. Cells were harvested and RNA was isolated using the RNeasy Micro Kit (Qiagen). The quality and quantity of the RNA samples was measured using the Agilent Fragment Analyzer 5300 system and Qubit (Invitrogen) respectively. 100ng of total RNA was used to prepare TruSeq Stranded mRNA libraries (20020594) following the manufacturer’s protocol with custom 384 xGen UDI-UMI adapters (IDT). Prepared libraries were validated with the Fragment Analyzer system dsDNA 910 Reagent Kit (35-1500bp) and Qubit dsDNA HS Assay Kit (Cat. Q32854). All libraries were then pooled in equimolar ratio and sequenced on a Nextseq2000 (Illumina) by using a P2 flow cell with 50bp paired-end reads.

### RNA-sequencing data analysis

Quality control on the sequence reads from the raw FASTQ files was performed with FastQC (v0.11.8). TrimGalore (v0.6.5) was used to trim reads based on quality and adapter presence after which FastQC was again used to check the resulting quality. rRNA reads were filtered out using SortMeRNA (v4.3.3) followed by alignment to the reference genome fasta (Drosophila_melanogaster. BDGP6.32) using the STAR (v2.7.3a) aligner. Followup QC on the mapped (bam) files was performed using Sambamba (v0.7.0), RSeQC (v3.0.1) and PreSeq (v2.0.3). Readcounts were then generated using the Subread FeatureCounts module (v2.0.0) with the Drosophila_melanogaster.BDGP6.32.105.chr.gtf file as annotation. CPM and RPKM Normalized versions of the counts table were generated using the R-packages edgeR (v3.28) as well as a normalized version using DEseq2 (v1.28). The differential expression analysis was performed using DESeq2.

### Statistics and graphs

All statistical analysis were performed in GraphPad Prism. Plots were generated in Python using Panda, Seaborn and matplotlib. Fig. 5E and chromosome icon in Fig. 1A were created with Biorender.com.

## Acknowledgements

We thank all the members of the Lens and Janssen lab for their valuable input during laboratory meetings. This work was funded by the European Research Council (ERC) under the European Union’s Horizon 2020 research and innovation program, grant agreement No. 850405, and VIDI VI.Vidi.203.001 financed by the Dutch Research Council (NWO). We acknowledge the Utrecht Sequencing Facility (USEQ) for providing sequencing service and data. USEQ is subsidized by the University Medical Center Utrecht and The Netherlands X-omics Initiative (NWO project 184.034.019).

