## Supplementary_figures for "DNA Double-Strand Break Movement in Heterochromatin Depends on the Histone Acetyltransferase dGcn5"

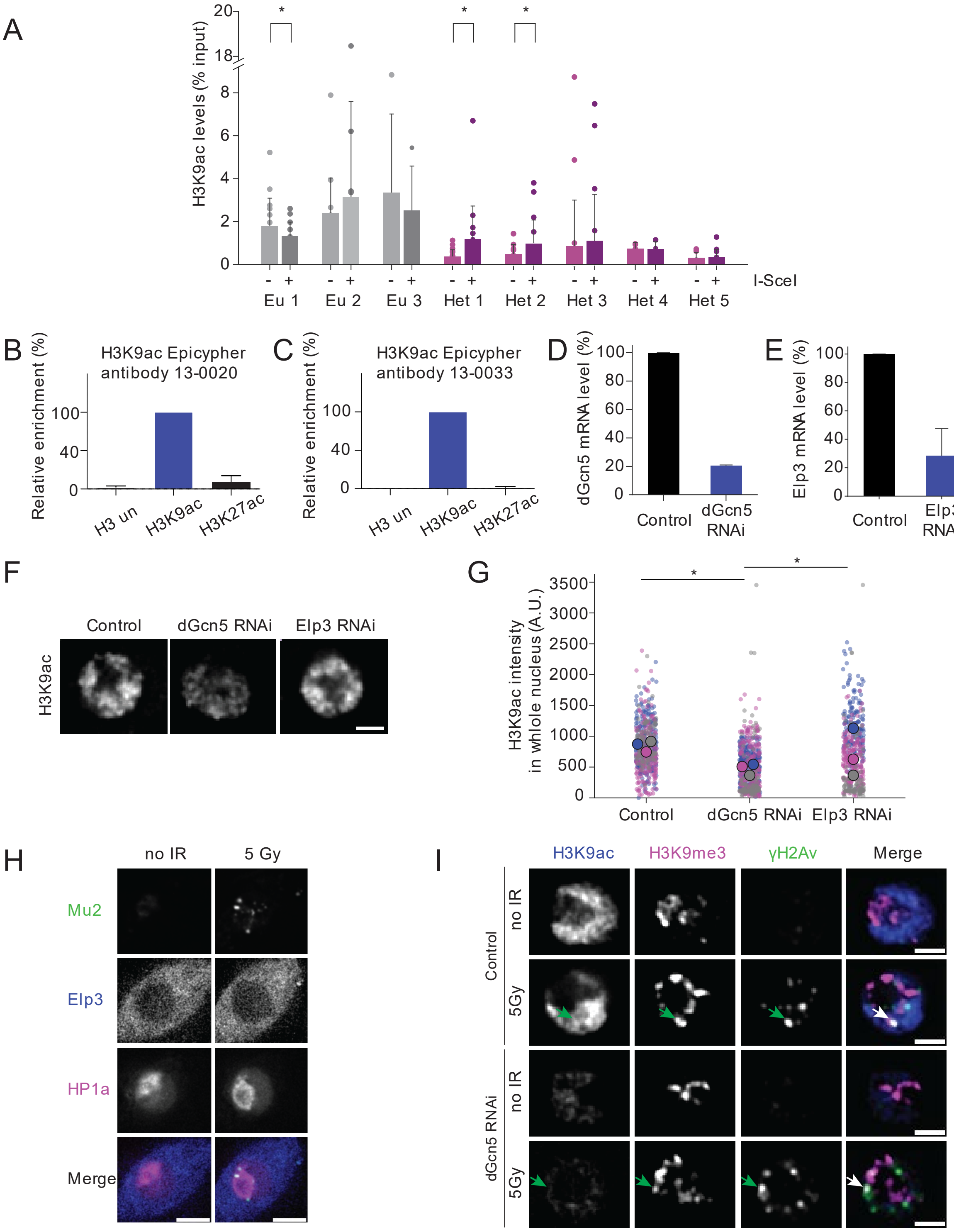


**Figure S1. dGcn5 controls H3K9ac levels in *Drosophila.*** **A**. ChIP-qPCR result for H3K9ac in three euchromatic and five heterochromatic DR-*white* lines, with (hsp.I-SceI) and without (no hsp.I-SceI) DSB induction. Graph indicates the percentage of input (normalized to internal *yellow* control gene) of H3K9ac from the same experiments as in *Fig. 1C*. Error bars represent mean +SD from ≥4 independent experiments. **B, C**. Specificity test for two anti-H3K9ac antibodies from Epicypher (*B*, lot number 13-0020, *C*, lot number 13-0033). Specificity was determined by using the SNAP-ChIP K-AcylStat nucleosome panel from Epicypher.

Error bars indicate mean +SD from ≥3 independent experiments. H3 un = unmodified H3. **D, E**. Normalized expression levels of dGcn5 (*D*) and Elp3 (*E*) in control (yellow dsRNA) and HAT-depleted Kc cells, determined by Reverse Transcription followed by quantitative PCR (RT-qPCR). Error bars represent mean +SD from 3 independent experiments. **F**. Immunofluorescence staining for H3K9ac in control, dGcn5-depleted and Elp3-depleted S2 cells. **G**. Quantification of absolute nuclear H3K9ac intensity levels in control, dGcn5-depleted and Elp3-depleted S2 cells. Each bigger circle (grey, blue, magenta) represents the average intensity within one experiment. Smaller circles represent individual cells within one experiment. **H**. Representative time-lapse images of non-irradiated and 5Gy-irradiated Kc cells of fluorescently tagged Mu2 (green, DSB marker), Elp3 (blue) and HP1a (magenta, heterochromatin marker). Images were taken 10 minutes after IR. Scale bars = 5µm. **I**. Representative immunofluorescence images for the quantification shown in *Fig. 1E, F*. Non-irradiated and irradiated control (yellow dsRNA) and dGcn5-depleted Kc cells were stained for H3K9ac (blue), H3K9me3 (magenta, heterochromatin marker) and γH2Av (green, DSB marker). Arrowheads indicate γH2Av foci within heterochromatin. For **F** and **I**, scale bars = 2µm. For **A** and **G**, (*) P-value≤0.05, paired t-test (*A*) and one way ANOVA followed by Tukey's multiple comparison (*G*). If not shown, P-value not significant (>0.05).

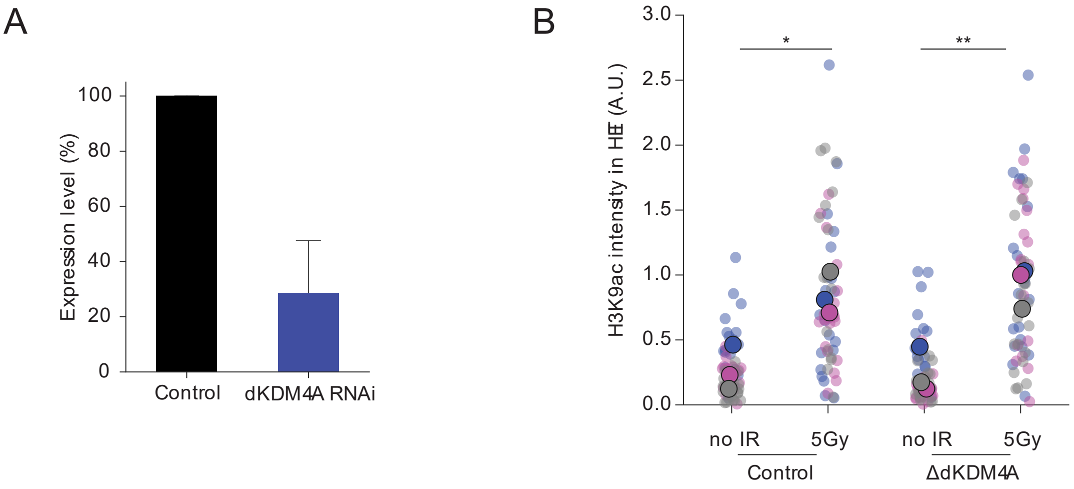


**Figure S2. Deposition of new H3K9ac marks at heterochromatic DSBs is independent of dKDM4A. A**. Normalized expression levels of dKDM4A in control (yellow dsRNA) and dKDM4A-depleted Kc cells, determined by RT-qPCR. Error bars represent mean +SD from 3 independent experiments. **B**. Quantification of H3K9ac intensity levels at DSBs in heterochromatin (DAPI bright) in non-irradiated versus irradiated control and ΔdKDM4A larval wing disc cells. For non-irradiated cells, H3K9ac signal was quantified in the entire DAPI bright (heterochromatin) area. Irradiated cells were fixed 5-10 minutes after damage induction. The graph represents 3 independent experiments per condition. Each big circle (grey, blue, magenta) represents the average intensity within one experiment. Small circles represent individual cells within one experiment. (*) P-value≤0.05, (**) P-value ≤ 0.01, one way ANOVA followed by Tukey's multiple comparison. If not shown, P-value not significant (>0.05).

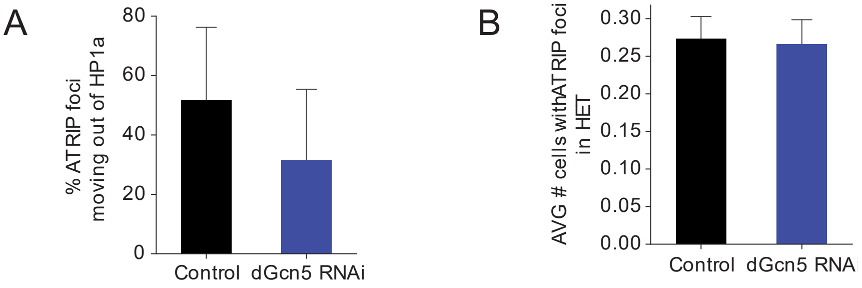


**Figure S3. dGcn5 promotes ATRIP foci movement outside heterochromatin. A**. Quantification of the percentage of ATRIP foci that eventually move out of HP1a domain as depicted in *Fig. 3B*. **B**. Quantification of the average number of cells with ATRIP foci inside HP1a domain upon control (yellow dsRNA) or dGcn5 depletion within the window of time imaged (120 minutes). Error bars represent mean +SD from 3 independent experiments. If not shown, P-value not significant (>0.05).

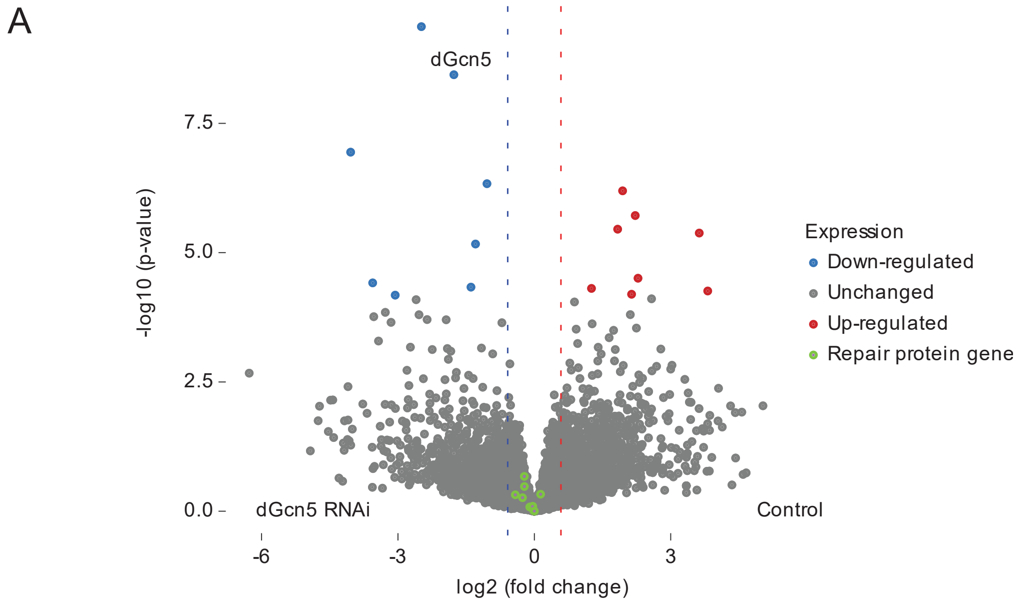


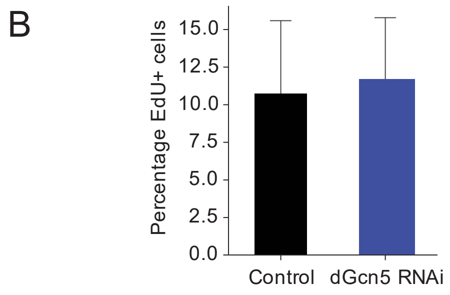


**Figure S4. dGcn5 knockdown does not affect cell cycle stage or transcription of DSB repair genes. A**. Volcano plot for bulk RNA-sequencing of Kc cells depleted of dGcn5 or yellow (control). See Table S1-3 for more details. **B**. Quantification of the percentage of EdU positive (S-phase) Kc cells upon dGcn5 and yellow (control) knockdown. Error bars represent mean +SD from 3 independent experiments.

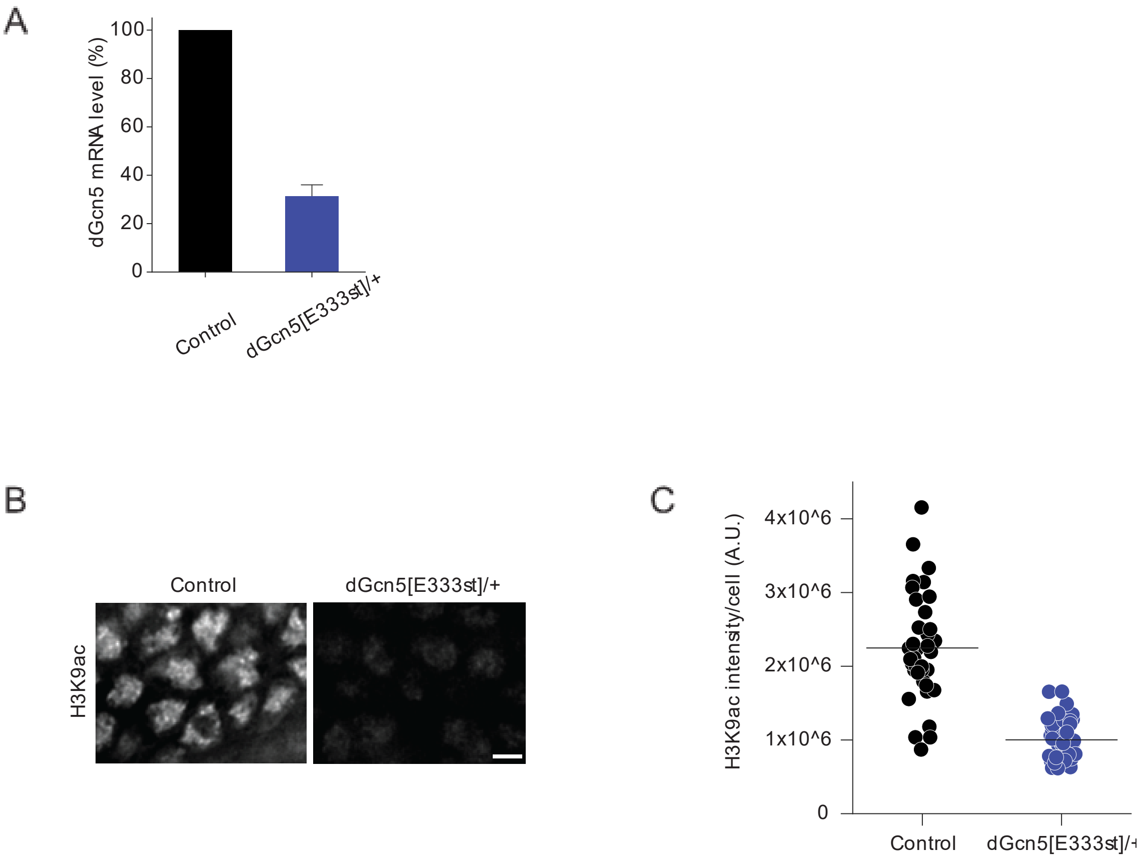


**Figure S5. dGcn5[E333]/+ mutant larval tissue has reduced H3K9ac nuclear levels. A**. Normalized expression levels of dGcn5 in control (OregonR) and dGcn5[E333st]/+ third instar larval tissues as determined by RT-qPCR. Error bar indicates mean +SD from 3 independent experiments. **B**. Representative images of *Drosophila* wing discs dissected from third instar larvae of OregonR (Control) or dGcn5[E333st]/+ and immuno-stained for H3K9ac. Scale bar = 2µm. **C**. Quantification of H3K9ac intensity per cell for the wing discs depicted in *B*. Lines represent mean intensity.
